# pyQCM-BraTaDio: A tool for visualization, data mining, and modelling of Quartz crystal microbalance with dissipation data

**DOI:** 10.1101/2023.12.15.571789

**Authors:** Brandon M. Pardi, Syeda Tajin Ahmed, Silvia Jonguitud Flores, Warren Flores, Laura L.E. Mears, Bernardo Yáñez Soto, Roberto C. Andresen Eguiluz

## Abstract

Here, we present a Python based software that allows for the rapid visualization, data mining, and basic model applications of quartz crystal microbalance with dissipation data. Our implementation begins with a Tkinter GUI to prompt the user for all required information, such as file name/location, selection of baseline time, and overtones for visualization (with customization capabilities). These inputs are then fed to a workflow that will use the baseline time to scrub and temporally shift data using the Pandas and Numpy libraries and carry out the plot options for visualization. The last stage consists of an interactive plot, that presents the data and allows the user to select ranges in MatPlotLib-generated panels, followed by application of data models, including Sauerbrey, thin films in liquid, among others, that are carried out with NumPy and SciPy. The implementation of this software allows for simple and expedited data analysis, *in lieu* of time consuming and labor-intensive spreadsheet analysis.

**Metadata:** 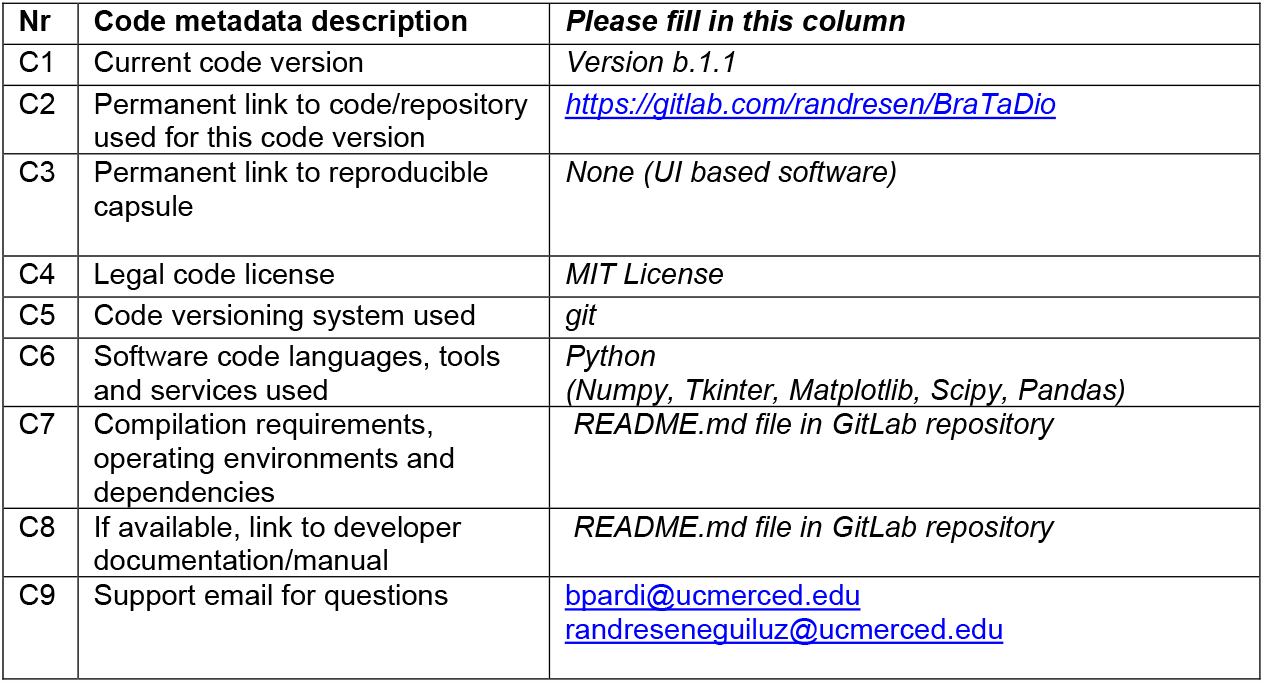

## 1. Motivation and significance

Quartz crystal microbalance with dissipation (QCM-D) is an acoustic-based, surface sensitive technique that measures small mass changes and energy losses, in real time, at the surface of a sensor. The sensors are commonly fabricated from a piezoelectric material, such as quartz, which allows thickness-shear of the sensors at its resonance frequencies by applying an alternating electric field. The most common cut of quartz crystals for QCM-D applications is the AT-cut, as it has excellent frequency-temperature characteristics.^1^ For these sensors, there are *n* number of acoustic modes, referred to here as overtones or overtone order, that can be approximated as standing waves perpendicular to the crystal surface with negligible longitudinal wave propagation.

The penetration depth of a 5 MHz shear wave in water is approximately δ ≈ 250 nm for the fundamental overtone and δ ≈ 70 nm for the 13^th^ overtone,^2,3^ making QCM-D a surface-sensitive instrument. For small films formed by analytes at the surface of the sensor, the resonance frequency *f* is inversely proportional to the total thickness of the plate. That is, the effective thickness *d*_eff_ = *d*_Q_ + *d*_f_ of the sensor increases with increasing amount of analyte coupled to the sensor surface, decreasing its resonance frequency *f*. This relationship was first identified by Sauerbrey,^4^ and is schematized in Figure 1. When the forming film is rigid or very thin (and homogeneous and continuous), the film-sensor interface does not change the bandwidth (Δ*Γ* = 0). However, if the forming films are viscoelastic (or heterogeneous or discrete), the shift or change in frequency Δ*f* is coupled with a change in bandwidth as well (Δ*Γ* ≠ 0), and the ideal inverse linear relationship between changes in thickness and changes in frequency no longer apply.^5^

**Figure 1.**
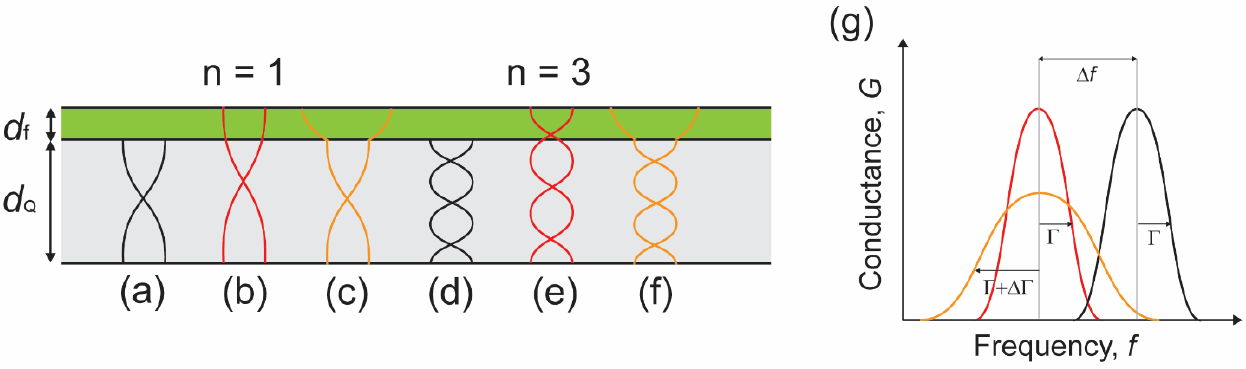
(a) and (d) Standing wave of overtone orders *n* = 1 and *n* = 3 across the thickness of the sensor, *d*_Q_. (b) and (e) Increased sensor thickness due to analyte film decreases resonance frequency. (c) and (f) When the analyte film is softer than the sensor, the interface generates a distortion to the standing wave, and the frequency shift becomes proportional to the mass, rather than the thickness. (g) Frequency and bandwidth shifts, Δ*f* and Δ*Γ*, respectively, are used to measure changes in film thickness *d*_f_ and energy losses.

QCM-D has gained popularity in many different scientific fields due to its experimental simplicity and versatility. QCM-D (or just QCM if not quantifying energy losses) can be combined with a variety of instruments for *in situ* complementary measurements, such as atomic force microscopy (AFM),^6^ microtribometry,^7^ surface plasmon resonance (SPR),^8^ or electrochemistry,^9^ among others. However, one drawback rests in linking experimental conditions, understanding relationships between changes in frequency Δ*f* and changes in bandwidth Δ*Γ*, to meaningful models. For example, films can be laterally homogeneous or heterogeneous, isotropic or anisotropic, the dissipation can take place inside the film or at a particle-surface region, continuous or discrete, and the bulk solution can be Newtonian or non-Newtonian. All these conditions will dictate the applicability of specific models. In a collection of works,^10–20^ Johannsmann and co-workers have developed, refined, and summarized qualitative and quantitative approaches for the analysis and interpretation of QCM-D data, which the reader is encouraged to review. Kanazawa and Gordon^21,22^ derived a simple relationship which expresses the change in frequency Δ*f* in contact with a fluid only in terms of the material parameters of the fluid (*i*.*e*., density *ρ*_F_ and viscosity *μ*_F_) and the quartz crystal (*i*.*e*., density *ρ*_Q_ and shear modulus *μ*_Q_). The relationship is valid for semi-infinite viscoelastic media. Du and Johannsmann^12^ derived the elastic compliance 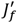of a viscoelastic film deposited on a quartz crystal surface in a liquid environment from the ratio of bandwidth shift Δ*Γ* and frequency shift Δ*f*. Voinova *et al*. derived the general solution describing the dynamics of two-layer viscoelastic materials of arbitrary thickness deposited on a solid (the quartz crystal) surface in a fluid environment.^23^ This relationship can be used for QCM-D measurements of layered structures, such as protein films adsorbed from solution (the analyte) onto the surface or swelling and deswelling of polymer brushes. Despite the relative simplicity with which QCM-D operates and its versatility, it becomes clear that quantitative results are non-trivial to obtain and require careful analysis.

In any QCM-D experiment, real-time monitoring of sensor surface-environment generates large volumes of data entries. Packages used to collect data do not typically possess straightforward data visualization, data mining capabilities, and basic model applications. Furthermore, programs associated with QCM-D data collection and analysis are often proprietary with limited access. There are other open source packages, such as RheoQCM^24^ and pyQTM,^25^ however, they focus on data modelling rather than data mining. Here, we present an intuitive Python-based, opensource software that is QCM-D manufacturer agnostic of multi-harmonic collecting systems for (1) simple and fast data visualization and interaction, (2) data mining and reduction, (3) crystal thickness calibration, and (4) basic model applications. The supported models include (i) Sauerbrey, for rigid thin films, (ii) Johannsmann, for viscoelastic thin film in a Newtonian liquid, and (iii) quartz crystal thickness determination. Lastly, we validate the software by visualizing and applying models to data collected with two commercial multi-harmonic QCM-D systems.

## 2. Software description

To perform the various levels of data visualization, applying models, and fitting possible with pyQCM-BraTaDio tool, it is crucial for the input data files to have the correct data structure and file extensions, depending on the source system. Supported data structures at the time of writing include QSense, QCM-I, and openQCM-D Next. Raw data is required to be exported to one of the following formats: *.txt, *.csv, *.xls, *.xlsx, and *.xlsm. The format structure for the data that each system generates, and more importantly, format structure that the pyQCM-BraTaDio tool converts to for further operations are detailed in the SI Input Data Structure section.

### 2.1. Software architecture

pyQCM-BraTaDio is a Python package implemented in Python 3.10.5, intended to be used as a data visualization (plotting), experimental data mining, and basic model application tool for multiharmonic QCM-D experimental data. It operates by following the workflow and file database organization shown in Figure 2. The main steps of the workflow are the generation of multiple data frames from the input file, customization of the plotting properties, selection of overtone orders (*n*) for display and analysis, selection and labelling of data ranges, calculation of average frequencies (*f*_n_) and dissipations (*D*_n_), average change in frequencies (Δ*f*_n_) and average change in dissipations (Δ*D*_n_) of selected and labelled data ranges, with corresponding standard deviations, and model (*e*.*g*., Sauerbrey mass, *m*_Sauerbrey_, sheardependent compliance of thin films, 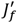, and quartz crystal thickness estimation, *h*_q_) selection. Selection of multiple data ranges is possible, for example, for selecting ranges corresponding to double-layers or buffer washes.

**Figure 2.**
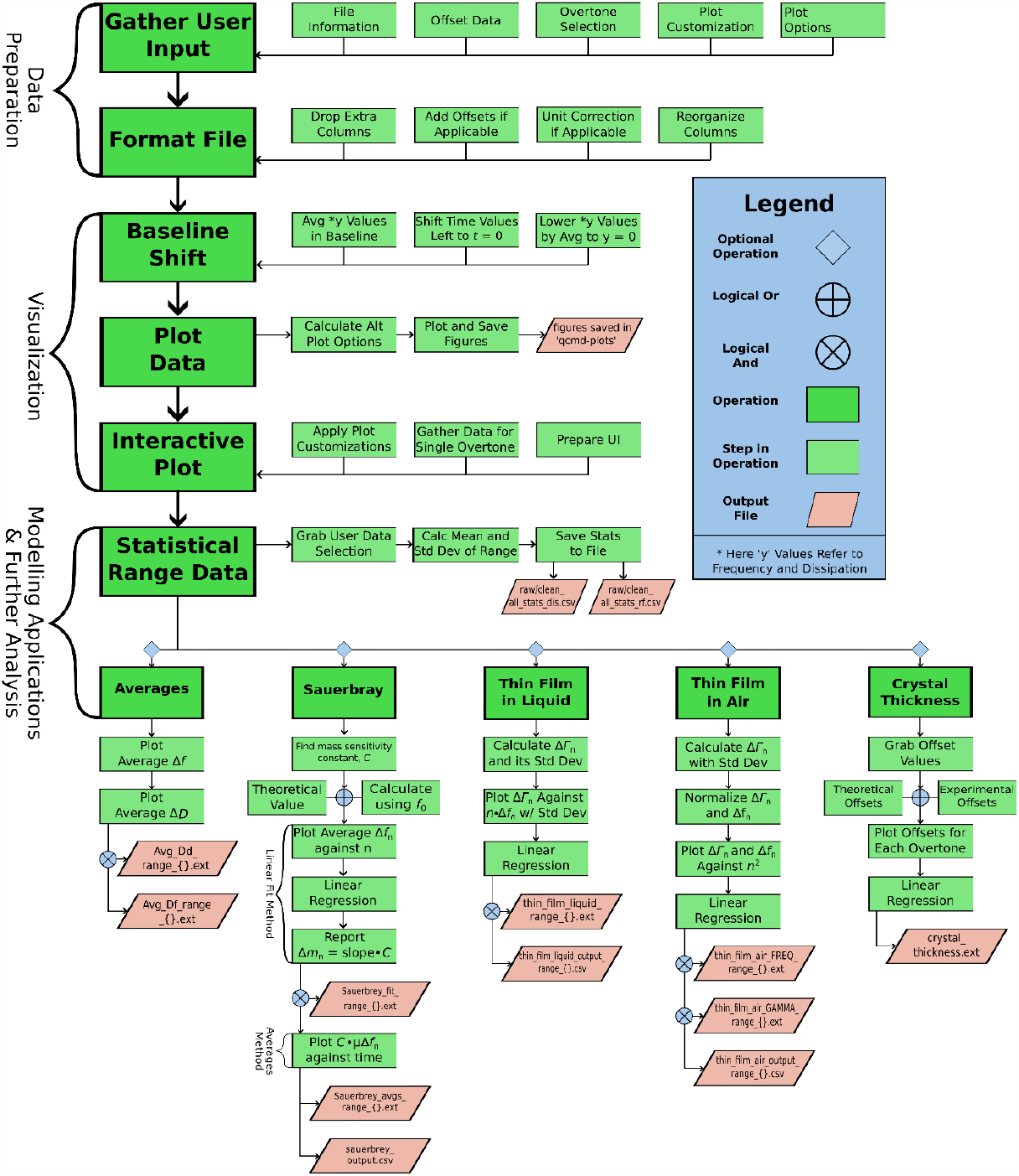
pyQCM-BraTaDio workflow and structure of operations. Beginning with data preparation, pyQCM-BraTaDio takes in the input fields specified by the user in the UI, and formats the file dependent on the experimental apparatus used to record the data. Once data is prepared, it is then visualized. This includes shifting data by the baseline, computing alternate plot options, saving figures, and generating the interactive plot. The optional modelling and further analysis layer can commence after visualization. This starts with the user making selections in the interactive plot and basic statistical calculations being applied on those selections, and ends with the execution of various available models for the data previously selected.

pyQCM-BraTaDio generates several output files. All generated plots are saved in the “qcmd-plots” directory and model application plots saved in a subdirectory within named “modelling” with the selected output format (*e*.*g*., PNG, TIFF, or PDF). Calculated averages and standard deviations of selected ranges using the interactive plot are saved in a *.csv file for the chosen overtone order frequencies and dissipations in the “selected_ranges” directory. pyQCM-BraTaDio includes python functions to read, compare, and statistically analyze the resulting *.csv files, detailed in the “*Comparison of experimental data to theoretical models*” section. Alternatively, the resulting files can be opened with standard text editors or data analysis tools such as Excel or Origin for further analysis.

pyQCM-BraTaDio utilizes several well-established open-source python packages. Most notably, Pandas,^26^ Numpy,^27^ for a variety of computational tasks, Tkinter for the interactive plotting engine and application user interface (UI),^28^ Matplotlib^29^ for the display of scatter plots and models and interactive plot interface, and SciPy^30^ for much of the model application capabilities. Pandas is a package providing flexible data structures to make working with relational or labelled data easy. It provides an intuitive way of working with data from spreadsheets, interacting with it with code. It is a high level, fundamental building block data for analysis in Python. NumPy is another core building block, as all the other libraries used rely on it as well. NumPy is used in almost every field of science and engineering in Python, containing multidimensional array and matrix data structures, allowing for easy operations on complexly related data. The number of available operations is enormous, and they are also guaranteed an optimized runtime. Matplotlib is the core visualization tool used for this software. It is a comprehensive library for designing and efficiently plotting static, animated, and interactive plots in Python. SciPy is the final pivotal library for this software. It provides highly optimized fundamental algorithms for model applications and analysis, such as integration, interpolation, statistics, and more. SciPy is the library behind our model application routines. It is a very powerful tool that is relatively simple to use in conjunction with our other libraries.

Our implementation begins with a Tkinter GUI to prompt the user for all required and optional information, such as file name/location, baseline time, which overtones to plot, and any additional data plot options. These inputs are then fed to an analysis workflow that will use the baseline time to scrub data using the Pandas library and carry out the plot options. The platform, most importantly, incorporates an interactive plot, that allows the user to choose the range of experimental data over time to calculate data averages and standard deviations. These will be used as input for the various model applications.

The implementation of this software allows for expedited data mining and analysis, *in lieu* of time consuming and laboring spreadsheet analysis needed for one experiment.

### 2.2. Software functionalities

The interaction with pyQCM-BraTaDio is via a GUI, which allows the user to interact with the software with minimal to no console interaction. The main window is organized into four main regions, shown in Figure 3. These regions are (1) initialization conditions, (2) selection of frequencies and dissipation for data mining, visualization, and model application purposes, (3) interactive plots for data mining, and (4) selection of plotting options and models.

**Figure 3.**
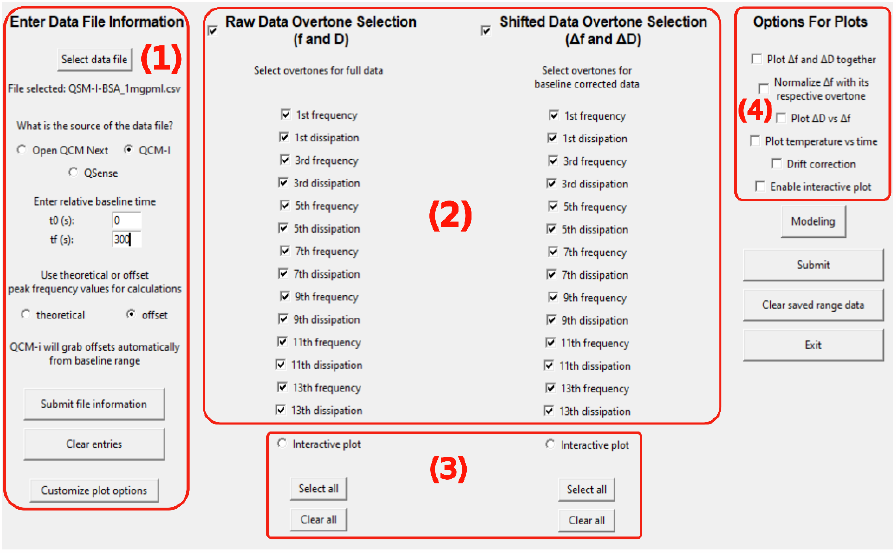
User interface of pyQCM-BraTaDio. (1) Initialization conditions, (2) selection of frequencies and dissipation for data mining, visualization, and modeling, (3) interactive plotting options for data range selection, and (4) selection of plotting options and modeling.

#### 2.2.1 Entering data file information

In region (1) from the main GUI shown in Figure 3, the user loads the relevant file in any of the supported formats: *.txt, *.csv, *.xls, or *.xlsx, and indicates the data structure by selecting the multi-harmonic instrument that generated the data. Next, the ‘*relative baseline time’* (*‘absolute baseline time’* for openQCM Next) refers to the beginning (*t*_0_) and end (*t*_f_) of what will be considered the experimental baseline. For example, if the file contains an air-to-liquid transition, followed by an equilibration time, the relative baseline time will consist of the last minutes of the equilibration time. pyQCM-BraTaDio calculates the average Δ*f*_n_ and average Δ*D*_n_ of all datapoints within the selected range (between *t*_0_ and *t*_f_) and uses those averages to set the reference (i.e., Δ*f*_n_ ≈ 0 Hz and Δ*D*_n_ ≈ 0).

The user is next asked to indicate usage of either theoretical or experimental offset values for resonant frequency calculations. Selecting theoretical uses resonant frequency values for 5 MHz AT-cut quartz crystals outlined in SI Table 1, selecting offset will lead to two potential workflows. If the user has selected QCM-I, the offset values are automatically taken from the user-inputted baseline time frame. If QSense is selected, the user will be prompted to either enter values in the window shown in Figure 4 or edit the ‘COPY_PASTE_OFFSET_VALUES_HERE.csv’ file in the ‘offset_data’ directory. It should be noted that these offset values are not required for basic visualization purposes, but model application functions will not be viable as they rely on the resonant frequency values for calculation. It is necessary to keep in mind that the typically reported values of dissipation *D*_n_ are related to bandwidth Γ_*n*_ by:^22^

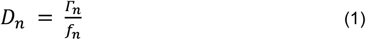

where *D*_n_ is the dissipation of order *n*, Γ_*n*_ is the bandwidth of order *n*, and *f*_n_ is the resonance frequency of order *n*.

**Figure 4.**
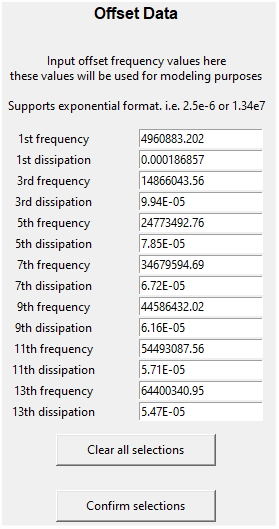
Input window for calibration obtained frequency and dissipation values (sometimes referred to as offset values).

Finally, it is necessary to submit input by clicking the ‘*Submit file information’*. The user also has the capability of personalizing the plotting aesthetics, such as size and font type for axis titles, *x* and *y* plotting ranges, selecting colors for data, among others. These options are described in the Customize Plot Options section in the SI.

#### 2.2.2 Data selection for visualization, mining, and analysis

The data selection region allows the users to choose the frequencies and dissipations from specific overtones for data mining, visualization, and model application purposes. pyQCM-BraTaDio works with overtones *n* = 1, 3, 5, 7, 9, 11, 13. It is also here where the user selects to work with the raw, full data range (*i*.*e*., from the first to last data point acquired) or with the experimental data (*i*.*e*., from a predetermined data point to the last data point acquired) and taking the changes in frequencies (Δ*f*_n_) and dissipations (Δ*D*_n_).

#### 2.2.3 Interactive plots

To facilitate the procedure of data mining, that is, selection of frequency and dissipation ranges for time ranges of interest, it is possible to interact with the data via the interactive plots in two ways. First, the user is required to select the *Interactive plot* option under either *Plot raw data (f* and *D)* or *Plot shifted data (*Δ*f* and Δ*D)*. Next, it is necessary to designate one of the available overtones for data visualization (*e*.*g*., *n* = 3 in Figure 5) and a label to the data range (*e*.*g*., After_PBS in Figure 5). Note that even though one overtone is selected for visualization, operations done in the interactive plot will apply to all overtones selected in the GUI. Lastly, to enable the interactive plot, the user must check the box labelled ‘enable interactive plot’. The interactive plot will then display *f* and *D* or Δ*f* and Δ*D*, depending on the selection. Figure 6 shows the latter case.

**Figure 5.**
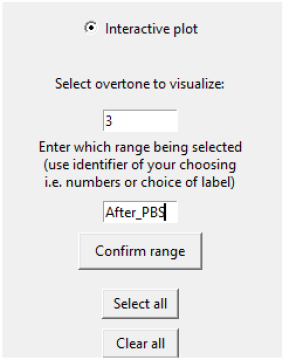
Activating the interactive plot option requires to select an overtone and assign a label or identifier for data mining.

**Figure 6.**
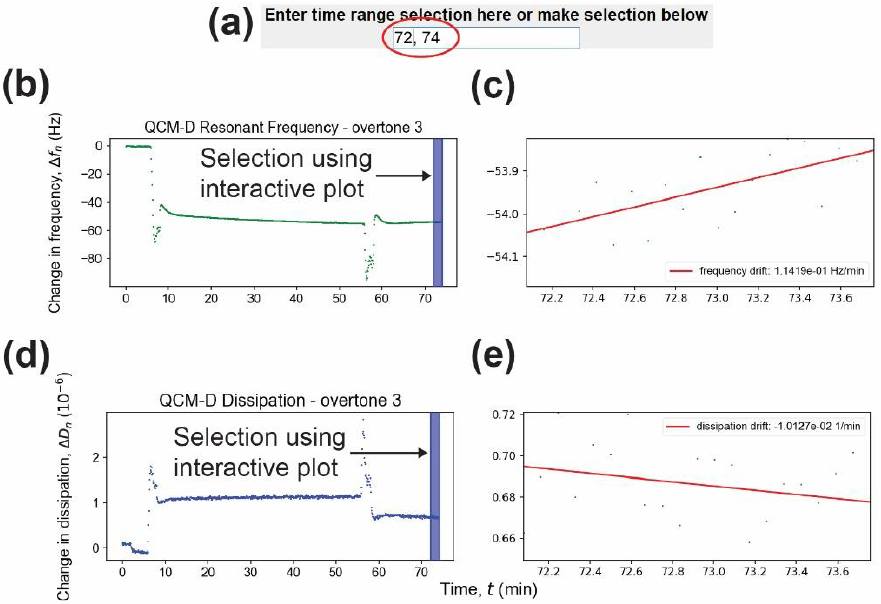
(a) Input line for time range selection, (b) change in frequency interactive plot, (c) zoomed-in region from the change in frequency interactive plot and frequency drift, (d) change in dissipation interactive plot, and (e) zoomed-in region from the change in dissipation interactive plot and dissipation drift.

The interactive plot window consists of 5 panels, an input text field to numerically provide a time range of interest in user-predefined units (default is seconds), Figure 6(a). The time range will be highlighted in the frequency, Figure 6(b), and dissipation, Figure 6(d), interactive plots. Alternatively, the user can select a range by clicking and holding the left mouse button in either direction of the frequency or dissipation interactive plots. The input text field will be updated and display the time limits of the selection. Interactive frequency and interactive dissipation plots are coupled, that is, same time ranges will be applied to frequency and dissipation channels. The plots shown in Figures 6(c) and (e) display the selected range only and calculated drift by applying a linear regression. For frequency, it will be in units of Hz over time, and numerical value over time for the dissipation (*i*.*e*., seconds, minutes, or hours, depending on the selected unit under the *Customize plot options* menu). pyQCM-BraTaDio will compute the average and standard deviation of the data points contained within the selected range for each overtone selected, every time a selection is made. The range can be moved and adjusted as needed, and the new average and standard deviation computed will be updated each time. Repeating the process overwrites the previously saved average and standard deviation for that specific range identifier. The average and standard deviation will be calculated for all selected overtones (all *f*_n_, Δ*f*_n_, *D*_n_, and Δ*D*_n_) in a file contained in the *selected_ranges* directory. Details on the file structure can be found in the *Description of directories and output files* section of the Supporting Information.

Changing the range identifier signals to pyQCM-BraTaDio that a new experimental step has been selected and averages and standard deviations will be saved in new fields (instead of overwriting the previously stored values). Note that when a new range is selected, data for previous ranges will remain untouched and new selections will only update for the currently identified range. This data will remain untouched until the user clicks the ‘*Clear saved ranges*’ button in column 4, which is recommended when switching to new experimental data, or using different models.

It should be noted that depending on the data file size, combined with computational power, processing time can vary between a fraction of a second to a few seconds. Faster processing times were demonstrated using a Ryzen 9 5900X 12 core (24 threads) processor at 4.5GHz with 32 GB of RAM, while the lower end of software execution speed was using a more conventional computer, consisting of an Intel i7-10510U 4 core (8 thread) processor at 1.8 GHz and 16 GB of RAM.

## 3. Illustrative examples

### 3.1 Plot options

As in any experimentally obtained data, visual inspection is crucial for an initial assessment of baseline stability, anomalies, such as presence of undesired air bubbles, leaks, signal loss, among others. Two basic visualization options are implemented in the pyQCM-BraTaDio tool: (i) a full data range visualization referred here as raw data, Figure 7, and (ii) the experimental data, referred here as shifted data, Figure 8. For these options, the user can select the overtone order(s) to visualize the frequency and dissipation in various plotting formats. Figure 7(a) and 7(b) show the absolute frequency *f*_n_ and dissipation *D*_n_ as a function of time *t* for *n* = 1, 3, 5, 7, 9, 11, and 13 for a BSA solution absorbing to a gold substrate, as described in the SI under ‘Model experiments’ section using a QCM-I unit. SI Figures 4 and 5 show similar plots to Figures 7 and 8 for an identical experiment performed using QSense. When plotting the raw data (absolute frequency and dissipation values), only the *Plot raw data* option with at least one overtone must be selected.

**Figure 7.**
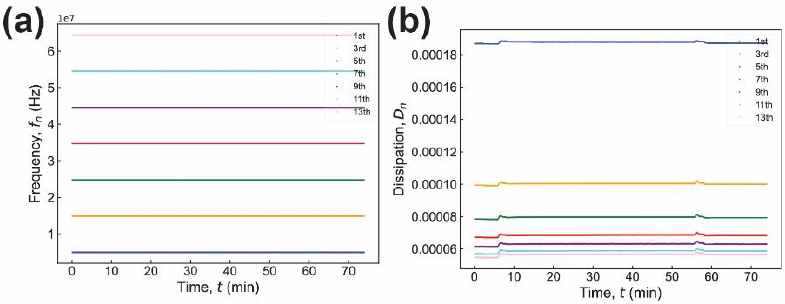
Raw data plots generated by BraTaDio for film formed from a solution of BSA at 1 mg/mL in PBS adsorbed to an Au-coated quartz crystal. (a) Absolute frequency *f*_n_ as a function of time *t*, (b) corresponding absolute dissipation *D*_n_ as a function of time for *n* = 3, 5, 7, 9, 11, and 13. The peaks seen in panel (b) correspond to transition periods, that is, pumping BSA after the PBS baseline was established (*t* = 5 min), and the second to a PBS wash (*t* = 55 min.

**Figure 8.**
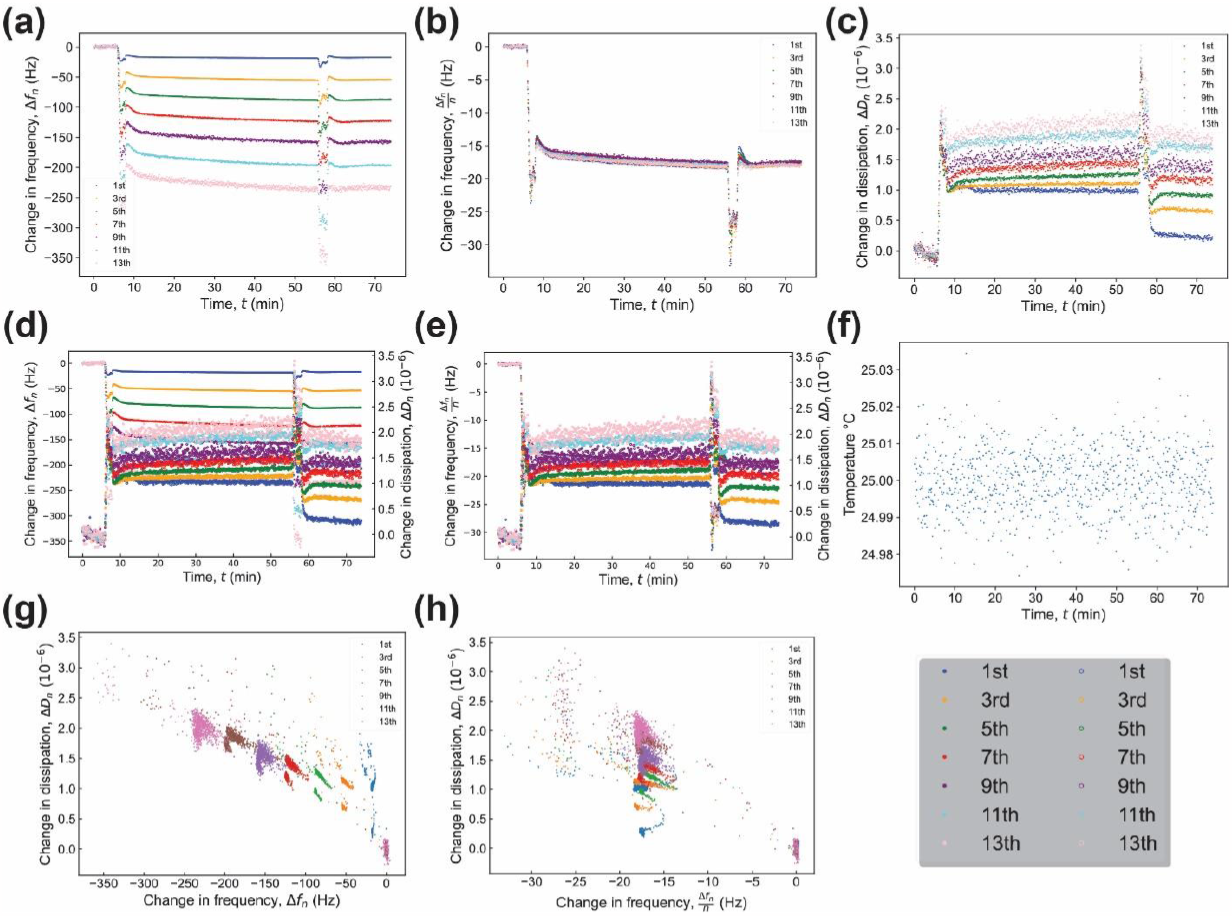
Plots generated by BraTaDio for a film formed from a solution of BSA at 1 mg/mL in PBS adsorbed to an Au-coated quartz crystal. (a) Change in frequency Δ*f*_n_ as a function of time *t*, (b) change in frequency normalized by overtone order, Δ*f*_n_ /*n* as a function of time *t*, (c) corresponding change in dissipation Δ*D*_n_ as a function of time, (d) change in frequency normalized by overtone order, Δ*f*_n_ /*n* and corresponding change in dissipation Δ*D*_n_ as a function of time for *n* = 5 and 7 for clarity, (e) change in dissipation Δ*D*_n_ as a function of change in frequency normalized by overtone order, Δ*f*_n_ /*n*, for *n* = 3, 5, and 7 for clarity, and (f) temperature T as a function of time. Data collected with a QCM-I system.

**Figure 9.**
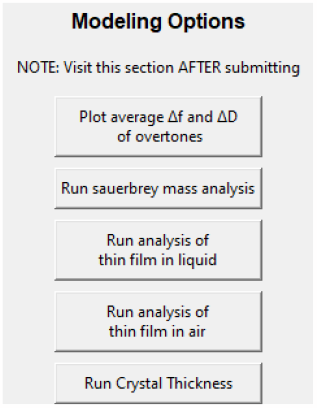
Available modelling options packaged in pyQCM-BraTaDio.

The relevant experimental data can be visualized by selecting the ‘*Plot shifted data*’ option. For example, change in frequency as a function of time, Δ*f*_n_ vs time *t*, Figure 8(a), change in normalized frequency as a function of time, Δ*f*_n_(/*n*) vs time *t*, Figure 8(b), change in dissipation Δ*D*_n_ vs time *t*, Figure 8(c), combined change in (normalized) frequency and change in dissipation as a function of time, Δ*f*_n_/*n* and Δ*D*_n_ vs time *t*, Figure 8(d), combined change in normalized frequency and change in dissipation as a function of time, Δ*f*_n_/n and Δ*D*_n_ vs time *t*, Figure 8(e), the temperature *T* as a function of time, *T* vs *t*, Figure 8(f), which is critical to determine any temperature effects in collected data. Finally, change in dissipation as a function of change in frequency, Δ*D*_n_ vs Δ*f*_n_(/n), Figure 8(g), and change in dissipation as a function of change in normalized frequency, Δ*D*_n_ vs Δ*f*_n_/n, Figure 8(h) to obtain qualitative insights of the adsorbed film rigidity. In these plots, commonly used for assessing differences in adsorbed protein^31–33^ or polymer films,^34^ a small slope is indicative of a compact and more rigid film, while a higher slope is indicative of a looser and more compliant layer. Change in frequency Δ*f*_n_ will always be plotted as normalized change in frequency Δ*f*_n_/n if the ‘*Normalize* Δ*f with its respective overtone*’ option is checked. The change in frequency Δ*f*_n_ and change in dissipation Δ*D*_n_ plots are automatically generated, and no selection is required. If one selects normalize Δ*f*_n_ with its respective overtone *n*, all associated Δ*f*_n_ plots will be normalized (Δ*f*_n_/*n*). This is not the case, however, for data used for model application purposes, as they will use normalized frequency if the model/equation requires it, regardless of the selection of this option. Note that when plotting shifted data (change in frequency and change in dissipation values), only the ‘*Plot shifted data’* option with at least one overtone must be selected.

### 3.2 Data analysis options

Matching QCM-D experimental data to models that provide physical interpretation is key for the quantitative characterization of liquids interacting with the quartz crystal surfaces or nanofilms. Models can be classified into steady-state models and kinetic models.^35,36^ With pyQCM-BraTaDio, it is possible to apply models of steady state (in equilibrium) thin films using one of the following models: (i) the Sauerbrey equation for very thin films,^4^ (ii) shear-dependent compliance of a thin viscoelastic film in a Newtonian liquid,^12^ and (iii) determination of the quartz crystal thickness.^37^ These models are described below, accompanied with experimental examples. It is important to note that the viscoelastic properties measured via QCM-D may deviate significantly from those obtained via bulk rheometers. That is in part, due to the fact that QCM-D measures at much higher frequencies (*i*.*e*., 5 – 65 MHz) than bulk rheometers (*i*.*e*., 0.1 – 100 kHz),^38^ and that the probing depth of QCM-D is of approximately 250 nm or less.^2,3^ For more complex models, the reader is referred to a review by Johannsmann *et al*.^11^

#### 3.2.1 The Sauerbrey equation for thin, rigid films

The Sauerbrey equation (eq 2) is a linear relationship between the resonance frequency change and the mass change of the acoustic oscillator:^4,10,11^

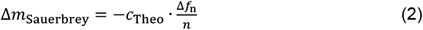

where Δ*m*_Sauerbrey_ is the change in Sauerbrey mass in (ng/cm^2^) (or mass per unit area), Δ*f*_n_ is the change in frequency of overtone *n* in (Hz), *n* is the overtone order, and *c*_Theo_ the mass sensitivity constant in (ng/cm^2^·Hz).

The Sauerbrey equation is applicable to very thin films, with a change in mass small compared to the quartz crystal, that can be considered rigid (no deformation) and perfectly coupled to the quartz crystal surface (no slip), homogeneous and evenly distributed over the quartz crystal surface. The theoretical value of *c*_Theo_ for a 5 (MHz) crystal is 17.7 (ng/(cm^2^·Hz)). The true *c*_True_ can be obtained from eq 3:

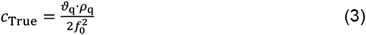

Where θ_q_ is the is the wave velocity in quartz, ρ_q_ is the density of quartz, and *f*_0_ is the measured, fundamental resonance frequency. It is frequent to report Δ*m*_Sauerbrey_ for one overtone order of interest, Figure 10(a). However, more accurate Δ*m*_Sauerbrey_ can be obtained from the slope of the change in frequency of each overtone Δ*f*_n_ as a function of overtone order *n* and multiplying it by the mass sensitivity constant, *c*_Theo_. Using the theoretical values is most of the times a good approximation, as *c*_Theo_ and *c*_True_ are usually in very good agreement. However, when the calibration values are provided, the Sauerbrey mass can be calculated using *c*_True_.

**Figure 10.**
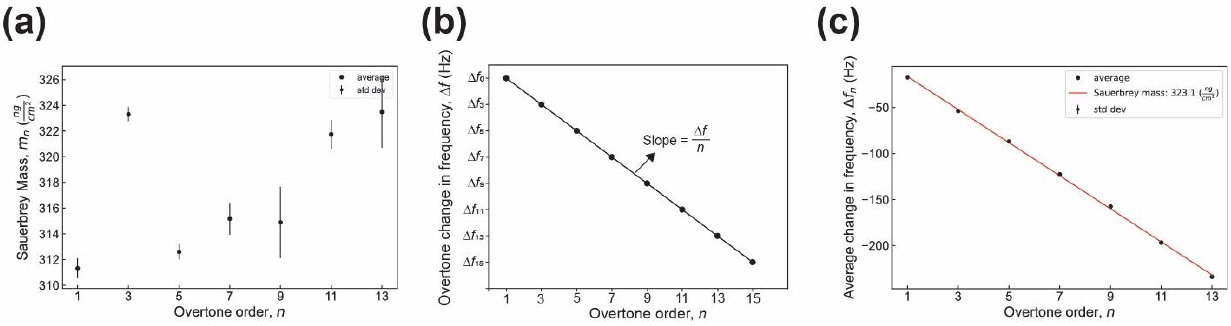
Sauerbrey mass as a function of overtone order. (a) Sauerbrey mass calculated as a funciton of each individual overtone order, (b) linear regression model to obtain the Sauerbrey mass using the changes in frequency from all available overtones, and (c) Sauerbrey mass calculated from performing a fit to the average change in frequency as a function of overtone order for the data range shown in Figrue 6 (BSA after PBS wash).

The data shown in Figure 10 corresponds to the Sauerbrey mass of a BSA film after a PBS wash formed from a 1 mg/mL bulk solution in PBS. Details are described in the *Model Experiments* section in the SI, and plots obtained from these experiments are shown in Figures 6, 7, and 8. pyQCM-BraTaDio plots the Sauerbrey mass for each selected overtone order, Figure 10(a) and saves the Sauerbrey mass values in *sauerbrey_output*.*csvs* file in the *selected_ranges_directory*. It also plots and displays the Sauerbrey mass obtained from calculating the slope of the linear regression and multiplying by the mass sensitivity constant *c*_True_, Figure 10(c). Similar results for an experiment conducted in a QSense system are shown in SI Figures 1, 4, and 5.

#### 3.2.2 Shear-dependent compliance of a thin viscoelastic film in a Newtonian liquid

For the cases in which the film formed on the surface of the quartz sensor is significantly more rigid than the environment Newtonian liquid, it is possible to obtain the shear-dependent compliance of the film by plotting the change in bandwidth over the negative of the change in frequency, 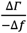, as a function of the overtone order, *n*, and calculating the slope:^11,12^

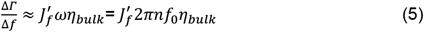

where 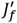is the shear dependent compliance of the film, ω the angular frequency (ω = 2*πn*), and *η*_*bulk*_ is the viscosity of the Newtonian fluid. This equation assumes that the viscous dependent compliance of the film is much smaller than the shear dependent viscous compliance of the bulk liquid, 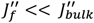. The model was derived by Du and Johannsmann and a detailed derivation can be found elsewhere.^12^

pyQCM-BraTaDio calculates the shear-dependent compliance of a thin viscoelastic film in a Newtonian liquid by computing the slope of the bandwidth shift for each overtone (ΔΓ_*n*_) as a function of the change in frequency times its overtone order value (*n* · Δ*f*_*n*_). Figure 11 shows the sheard-dependent compliance of a BSA film after a PBS wash formed from a 1 mg/mL bulk solution in PBS. The calculated value 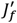is 0.0192 Pa^-1^. The datapoints displayed in the generated plot are saved in *thin_film_liquid_output*.*csv* file in the *selected_ranges* directory. Experimental details are described in the *Model Experiments* section, and plots obtained from these experiments are shown in Figures 6, 7, and 8.

**Figure 11.**
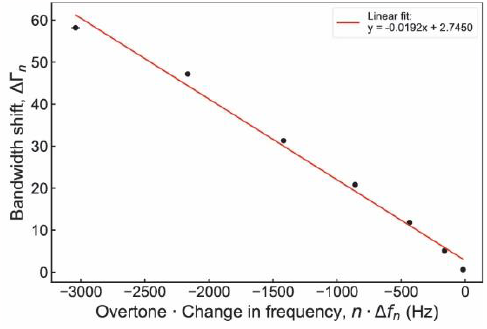
Shear dependent compliance, 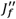of a BSA film formed from a bulk solution of 1 mg/mL BSA in PBS after a PBS wash.

**Figure 12.**
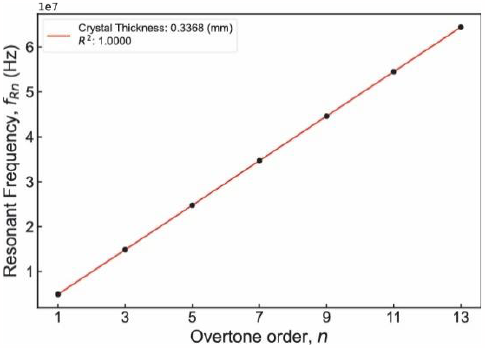
Determination of the quartz crystal thickness.

#### 3.2.3 Shear-dependent compliance of a thin viscoelastic film in air

For the cases in which the film on the surface of the quartz sensor is exposed to air as the medium instead of a liquid, the inertial effects of the medium become negligible. It is possible to obtain the shear-dependent compliance of the film by plotting the normalized change in frequency 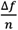, as a function of the square of the overtone order, *n*^2^, and calculating the slope:^11,24,25^

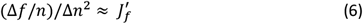

If the slope is relatively constant, it suggests that the shear-dependent compliance 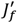is relatively insensitive to the frequency.

#### 3.2.4 Estimation of the quartz crystal thickness

For the cases in which the true thickness of the crystal is required, and the calibration resonance peaks known, it is possible to estimate the thickness of the crystal *h*_q_. The implementation is based on the work by Reviakine *et al*.,^37^

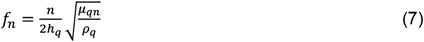

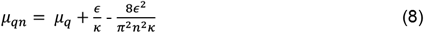

where *n* is the overtone order, *h*_q_ is the crystal thickness, *ρ*_q_ the density of quartz (2650 kg/m^3^), *μ*_qn_ is the elastic modulus of quartz considering piezoelectric stifening, *μ*_q_ is the shear modulus of quartz (2.93·10^10^ Pa), *ε* is the piezoelectric stress coefficient (-9.24·10^-2^ C/m^2^), and *κ* is the dielectric constant (3.982·10^-11^ F/m).

pyQCM-BraTaDio calculates the thickness of the quartz sensor for the cases in which the calibration resonance frequency values have been provided. The corresponding outputs are a plot displaying the calculated *h*_q_ from the user defined number of overtones and saves the computed value in the *crystal_thickness_output*.*csv* file in the *selected_ranges* directory.

## 4. Impact

QCM-D data model applications rely on the *a priori* knowledge of various quartz crystal properties, such as fundamental resonance frequency and bandwidth, *f*_0_ and Γ_0_ respectively, higher overtone order resonance frequencies and bandwidth, *f*_n_ and Γ_n_ respectively, and theoretical mass sensitivity constant *c*_Theo_ (-17.7 ng/cm^2^·Hz at 5 MHz). For those cases in which only changes in frequency Δ*f*_n_ and changes in dissipation ΔΓ_n_ without offset values (*e*.*g*., data coming from a QSense system) are exported or theoretical values are not acceptable for model application purposes, it is possible to read a calibration file containing the measured resonance frequencies and bandwidths of a free oscillating crystal in air and use those values to calculate, for example, the true mass sensitivity constant *c*_True_ or crystal thickness *h*_crystal_. Details on how to obtain calibration files from each compatible system with pyQCM-BraTaDio and how to generate/obtain a calibration are detailed in the Supporting Information.

## 5. Conclusions

pyQCM-BraTaDio is a Python software implemented *ad hoc* to expedite the process of data mining and analysis of QCM-D experimental data. We beging with a Tkinter GUI for metadata collection. These inputs and data are fed to several routines to mine and shift reformatted data with the Pandas and NumPy libraries. The user is able to interact with QCM-D in a novel way via a MatPlotLib interactive plot widget towards the end of the workflow. This interaction offers the user to apply several models such as Sauerbrey, thin film in liquid, thin film in air, and crystal thickness. This tool is key for efficient data analysis in preference over laborious spreadsheet evaluation.

## 6. Acknowledgements

R.C.A.E. acknowledges funding from the NSF-CREST: Center for Cellular and Biomolecular Machines through the support of the National Science Foundation (NSF) Grant No. NSF-HRD-1547848. R.C.A.E. and B.P. acknowledge funding from the CAREER grant NSF CMMI Grant No. #2239665 awarded to R.C.A.E.

## 7. Supporting Information

The authors have compiled additional supporting information in a separate document containing more details on the software’s execution, as well as demonstrating the efficacy of the software across multiple QCM-D devices.

