## Supporting Information for "pyQCM-BraTaDio: A tool for visualization, data mining, and modelling of Quartz crystal microbalance with dissipation data"

*Roberto C. Andresen Eguiluz

Science and Engineering 2, Room 292

Department of Materials Science and Engineering

University of California Merced

5200 N. Lake Rd., Merced CA 95340, USA

**Keywords:** quartz crystal microbalance with dissipation; analysis tool; open software.

### 1. Data interaction

#### 1.1 Input file structure

At the time of publication, pyQCM-BraTaDio supports 4 of the major QCM-D devices: openQCM-Next, QCM-I, QSense, and AWSensors. Below is a detailed description of the structure of the files these devices output, and how it is relevant to the software's execution, specifically the file formatting process.

The file structure of openQCM-Next is the file structure that inspired the structure of pyQCM-BraTaDio. Time here is recorded as absolute, meaning every entry has a time stamp in the format of hh:mm:ss, where as other formats record time as relative meaning time starts at 0 counts upwards from there. Columns are listed 'Frequency_n', 'Dissipation_n' where *n* = 0, 1, 2, 3, corresponding to overtones fundamental, $3^{rd}$, $5^{th}$, $7^{th}$, and $9^{th}$. The time column is labelled 'Time' in seconds, and temperature column labelled 'Temperature' in degrees Celsius.

QCM-I records time relatively and names its columns as 'Channel A Fundamental Frequency [Hz]', 'Channel A 3. Overtone [Hz]', 'Channel A 5. Overtone [Hz]', …, 'Channel A 13. Overtone [Hz]' and 'Channel A Fundamental Dissipation [ ]', 'Channel A 3. Dissipation [ ]', Channel A 5. Dissipation [ ]', …, 'Channel A 13. Dissipation [ ]'. Its time column is written as 'Channel A QCM Time [sec]' and temperature as 'Channel A Temp [Celsius]. These are the only necessary columns for pyQCM-BraTaDio's workflow. Others are dropped during file formatting. It should be noted that QCM-I also records frequency variation $\Delta f$, and dissipation variation, $\Delta D$, however we opt to use the full values, *f* and *D* respectively*.* This is due to some of the requirements for modelling downstream in the pipeline that rely on the full values, rather than deltas. Note that we refer to frequency variation and dissipation variation as change in frequency and change in dissipation, respectively.

Both openQCM-Next and QCM-I record data as *f* and *D*, so no offsets need to be considered. There is also no unit conversion required, as frequency and dissipation are recorded in their base units (*i.e.*, frequency values are recorded in Hz not MHz, and dissipation values are reported as $10^0$ and not $10^{-6}$ ).

QSense and AWSensors on the other hand, records $\Delta f$ and $\Delta D$, rather than *f* and *D.* QSense and AWSensors normalize their data by the overtone order ($\Delta f_n/n$), and report dissipation values as $10^{-6}$, meaning un-normalization and scaling is necessary. QSense and AWSensors data files require the most computation to preprocess, due to the three data operations described that other formats do not require. These operations are described in further detail in the next section.

#### 1.2 Working with files from QCM-D devices not natively supported by pyQCM-BraTaDio

If you have a device outside of the four supported devices mentioned above, consider the options below and reach out the authors with any questions or concerns. We are also able and willing to meet with readers and discuss adding their file types to be natively supported by pyQCM-BraTaDio.

##### 1.2.1 Manually formatting data files

The simplest approach is to manually format non-supported data files. However, it is time-consuming. To do so, follow the below steps:

1. Convert file to *.csv.

If the data file is in *.txt, *.xls, *.xlsx, etc, it must be converted to *.csv by saving the file as *.csv using Excel, or any other spreadsheet software.

2. Remove and rename columns.

Remove any columns that are not time, temperature, or the frequency/dissipation values for each overtone, as pyQCM-BraTaDio will not need these. pyQCM-BraTaDio references column names using the Pandas Python library. This means that the headers of your sheet must match the column names of the sample *.csv file provided in the 'raw_data' directory.

3. Data formatting.
   a. Magnitude scaling. QSense records dissipation as multiples of $10^{-6}$. pyQCM-BraTaDio reads dissipation as $10^{-0}$. Therefore, the user needs to ensure magnitudes are in terms of $10^{0}$ by multiplying all dissipation values by $10^{-6}$.
   b. Unnormalize data if normalized. pyQCM-BraTaDio relies on non-normalized frequency data (and dissipation if needed). If the user's experimental data is normalized, simply multiply each overtone's data by its corresponding overtone number.
   c. Offset value addition. For proper modeling purposes, change in frequency and change in dissipation columns need to be converted to absolute values. To do this, add offset data for each overtone, to its corresponding overtone's column in the dataset.

This process results in a file that pyQCM-BraTaDio can read. Please note, when saving this type of file following these formatting steps, prepend the word 'Formatted' to the data file (*i.e.*, if the data file was originally named 'qcmi_sample.csv', it should be renamed to 'Formatted-qcmi_sample.csv'). Additionally, if the file records time relatively (*i.e.*, starting at $t$ = 0 seconds) select the QCM-I option in the file options of the GUI. If the file records time absolutely (*i.e.*, using time stamps in the format of HH:MM:SS), then select openQCM-Next.

#### 1.2.2 Altering code to automate formatting

This option is more difficult to set and recommended for people with some degree of Python experience, but more efficient long term as it requires the user to add code to the 'format_file.py' file to automate the process described above. In that file, the function 'format_qsense' is a good example.

1. Define a new function as follows:

   def_format_<name of experimental device>(fmt_df, calibration_df)

Where <name of experimental device> is replaced with an identifier of the device used to generate the data, 'fmt_df' is the dataframe to be formatted, and 'calibration_df' is the dataframe containing offset data that will be added to 'fmt_df'. 'calibration_df' is only required if user's data is recorded as changes (*i.e.*, $\Delta f$ and $\Delta D$) rather than absolute values, as is the case with QSense. If the device records absolute values (*i.e.*, $f$ and $D$) then one can follow 'format_qcmi' as a more relevant example.

2. Define a dictionary called 'renamed_cols_dict' that will contain the key:value pairs, where the key is the original column name, the value is the new column name. For convenience, the new column names to be formatted are globally defined as a list at the top of the script. Using QCM-I formatting as an example, the dictionary will be defined as follows:

```
renamed_cols_dict = {'Channel A QCM Time [sec]':'Time',
        'Channel A Fundamental Frequency [Hz]':freqs[0],'Channel A Fundamental Dissipation [ ]':disps[0],
        'Channel A 3. Overtone [Hz]':freqs[1], 'Channel A 3. Dissipation  [ ]':disps[1],
…
'Channel A 13. Overtone [Hz]':freqs[6], 'Channel A 13. Dissipation  [ ]':disps[6],
        'Channel A Temp [Celsius]':'Temp'}
```

With the renaming dictionary defined, next is to call the ‘rename_cols’ function to rename the columns, passing the original dataframe and the dictionary defined. This function returns the formatted dataframe and does not format in place, therefore it is important to set a new variable, ‘fmt_df’ equal to this function. It should also be noted that this function drops columns that are not needed for pyQCM-BraTaDio’s execution.

It is at this point that if data is recorded in the format akin to QCM-I as described earlier, skip step 3.

3. Data formatting (magnitude order conversion, un-normalization, and offsets addition) using provided functions.

Note before proceeding, data may not need all formatting methods. Consult the device manufacturer to understand how the data is recorded, and proceed accordingly applying only needed formatting. That is, data may be recorded as full values, *f* and *D* but it may be in different units and/or be normalized, which as described above, needs correction.

Also note, the order for these operations is crucial, follow the order as described below.

a. Magnitude order conversion in pyQCM-BraTaDio is done via an applied lambda function. Navigate to the ‘format_qsense’ function in ‘format_file.py’ and copy the line:

```
fmt_df.loc[:, disps] = fmt_df.loc[:, disps].apply(lambda x: x*1e-6)
```

and paste this into the function, below the column formatting described in step 2. Note that this converts only magnitudes of dissipation values to multiples of $10^0$. Magnitudes may vary, so adjust accordingly.

b. Set the ‘fmt_df’ equal to the ‘unnormalize()’ function as follows:

```
fmt_df = unnormalize(fmt_df)
```

This will multiply all frequency values by their respective overtone number. Note that if dissipation is also normalized, it is required to adjust the ‘unnormalize()’ function accordingly.

c. Set the ‘fmt_df’ equal to the ‘add_offsets()’ functions as follows:

```
fmt_df = add_offsets(calibration_df, fmt_df)
```

This is where the calibration file is required. For each overtone it adds the offset value to all recorded values of that respective overtone in the data file.

Now return the fmt_df and proceed.

4. Add to the *if-elif* block in ‘format_raw_data.py’.

If the file does not need the offsets, follow the QCM-I formatting:

```
elif src_type == 'QCM-i':
    formatted_df = format_QCMi(data_df)
```

Copy/paste this above the last *else* statement in the *if-elif* block that handles the error case. Then replace the name of the function defined in step 2, and replace the string in quotes that is being evaluated against the ‘src_type’ variable, with a sensible device identifier. This will be referenced later in the GUI.

5. Adding the data file as an option to choose from in the UI.

Navigate to the ‘srcFileFrame’ class in ‘main.py’. Observe how the other options are added, and insert the file source type in the same fashion.

a. Add the file source type to the list:

self.file_src_types = ['QCM-d', 'QCM-i', 'Qsense', 'AWSensors', ‘NEW_FILE_SRC_TYPE_HERE’]

This should match exactly what you put to compare against ‘src_type’ in the previous step.

b. Add a radio button for the new file source option. You can copy paste the following lines:

self.opt3_radio = tk.Radiobutton(self, text="QSense ", variable=self.file_src_var, value=2, command=self.handle_radios)

self.opt3_radio.grid(row=2, column=0, columnspan=2)

Changing the 3 to a 4 in both instances of ‘self.opt3’, the string in text=’Qsense’ to the name of the experimental device, the number in ‘value=3’ to a 4, and number in ‘column=0’ to a 1

6. Adjust the ‘handle_radios’ function

In the same class as step 5, scroll down to the ‘handle_radios’ function. If data is recorded in absolute time, add the new source file ‘file_src_type’ (the string you appended to the list in step 5a) to the top if line as follows:

If self.file_src_type == ‘QCM-d’ or self.file_src_type == ‘NEW_FILE_SRC_TYPE_HERE’’:

If the data is recorded in relative time, add it to the next if line:

If self.file_src_type == ‘QCM-i’ or self.file_src_type == ‘NEW_FILE_SRC_TYPE_HERE’’:

After following these steps, proceed with data files as with any of the supported formats in pyQCM-BraTaDio.

An OpenQCM-D Next from Novaetech SRL (Pompei, Italy), controlled with openQCM-NEXT-0.1.2. A peristaltic pump (Golander LLC, BQ80S Microflow Variable-Speed) was used to inject buffers and molecule solutions. The native format of the experimental data is already in *.txt format.

File using the "add all" (to include all raw data collected) option before exporting, SI Figure 1(b). OpenQCM-Next generates a file containing all raw data in *.txt format and does not require any export operation, SI Figure (c).

**SI Table 1.** Theoretical resonance frequency values for 5 MHz, AT-cut quartz crystals.

| **Overtone order, *n*** | **Frequency, *f* (Hz)** |
|---|---|
| 0 or 1 | 4930000.00 |
| 3 | 14800000.00 |
| 5 | 24700000.00 |
| 7 | 34600000.00 |
| 9 | 44500000.00 |
| 11 | 54400000.00 |
| 13 | 64300000.00 |

#### 1.3 Description of directories and output files

pyQCM-BraTaDio generates many different output files during execution. Below is the list of directories and files that may be found in them.

- offset_data
    - contains the file 'COPY-PASTE_OFFSET_VALUES_HERE.csv'.
    - This is the file the user will copy and paste offset values if desired.
    - Alternatively, if user opts to use the offset data window to enter values, this is the file that those values are saved to.
- qcmd-plots
    - Contains ALL plots output by the software.
        - Note that different plotting options are indicated by the file name, but files will be overwritten if the same plots are done again even with different data.
    - modeling
        - Contains all plots generated from functions in the modeling window of the UI.
- raw_data
    - default directory for choosing the data the user would like to work with in the software.
    - Will contain sample data files upon first use, however any other files are placed here by the user.
- selected_ranges
    - contains output data from selections made in the interactive plot, as well as any modeling functions that utilize this data. These files include:

- clean_all_stats_dis.csv and clean_all_stats_rf.csv will contain the statistical data from selections made in the interactive plot of the baseline corrected data, as well as the user-specified name of the range, x-axis (time) bounds, and data file origin.
- Crystal_thickness_output.csv – overtone, offset values, curve fit of the offset values, and the crystal's thickness corresponding to each overtone.
- raw_all_stats_dis.csv and raw_all_stats_rf.csv are the same as the clean variant, just for the raw data.
- Sauerbrey_output.csv – contains overtone, average change in frequency with error, curve fit value of the average change in frequency, average Sauerbrey mass with error, quartz crystal constant calculated by the software, name of the range these values originate from, and the data file the data originates from
- Thin_film_air_output.csv
- Thin_film_liquid_output.csv – contains overtone order times the change in frequency for each overtone and its corresponding bandwidth shift, curve fit value of the bandwidth shift, and the range name and data source these values originate from

#### 1.4 Obtaining calibration files

Of the natively supported devices, openQCM Next, QCM-I, and QSense, QSense is the only device that generates data that will require an offset value file for more extensive analysis. The offsets are unnecessary if one only wishes to use basic visualization. However, more than just the deltas, $\Delta f$ and $\Delta D$, are required for any functionality beyond. openQCM Next and QCM-I record these values and do not need offset values. For QSense, after the experiment is completed, a *.QSD (or equivalent proprietary) file is generated. This file can only be accessed using QTools (or equivalent proprietary) software. Once opened, the data from the experiment is shown as:

$$Displayed\ frequency\ value = \frac{(real\ value - offset)}{overtone}$$

$$Displayed\ Dissipation\ value = \frac{(real\ value - offset)}{(1 \times 10^{-6})}$$

The file can be exported either as an *.txt or *.xls file. As mentioned earlier, in order to further analyze data using pyQCM-BraTaDio, the real frequency and dissipation value is needed. These values will be later entered either into the offset value window in the UI, or pasted directly into the 'offset_data/ COPY-PASTE_OFFSET_VALUES_HERE.csv' file

### 2 Model experiments

For the data visualization in the main text, we provide here the experimental details for how that data was obtained.

#### 2.1 QCM-D experiment

*Materials*

Phosphate Buffer Saline or PBS (Gibco, Catalog No: 10-010-031), Ethanol (ACROS, absolute, 200 Proof, ≥ 99.5%, Product # 61509-5000, Lot # B0542545A), Sodium dodecyl sulfate (MP

Biomedicals, ultra-pure, ≥99%, Catalog # 811032, lot # S0709) were purchased. For making base piranha solution: Ammonium Hydroxide (Chemsavers, Product # AMHE500ML, lot # AMHE080521, 28-30% pure), hydrogen peroxide (PERDROGENTM by Honeywell, MDL # MFCD00011333, ≥30% (w/w) stabilized) were used. Gold-coated silica quartz crystals were purchased from Quartz PRO (Product # QCM5140CrAu120-050-Q, resonance frequency- 5 MHz). Ultrapure water was collected from Thermo Fisher Millipore UV water purification system. Bovine Serum Albumin was purchased from Sigma Aldrich (Product # A3294) and prepared at 1 mg/ml in PBS at pH 7 to mimic physiologically relevant conditions.

*Methods*

Surface preparation for Quartz Crystal Microbalance with Dissipation (QCM-D)

The gold-coated crystals of 5 MHz base resonance frequency were rinsed copiously with ultrapure water (18.2 MΩ.cm), 2 wt% SDS, and ethanol, respectively, and repeated 3 times. After rinsing, the surfaces were dried with a stream of N2, cleaned with oxygen plasma, and stored in Petri dishes before use.

After using the crystals for protein adsorption experiments in the QCM-D, they were reused after cleaning in base piranha (6:1:1 by volume of water: ammonium hydroxide: hydrogen peroxide) by submerging the surfaces for ~20 seconds at 600 °C. Then the crystals were further rinsed with water and ethanol copiously several times, and then dried with $N_2$.

*Quartz Crystal Microbalance with Dissipation (QCM-D) setups*

QCM-D is a non-destructive acoustic shearing technique that uses a piezoelectric quartz sensor to measure changes in frequency and dissipation in real-time. The adsorption of molecules to the gold-coated crystal was assessed in a QCM-I by Gamry MicroVacuum Ltd. (Budapest, Hungary) or QSense (Biolin Scientific, Sweden). All measurements were at 25 °C. PBS was circulated through the system and allowed approximately 1 hr to equilibrate and establish the baseline. Then the BSA in PBS solution at 1 mg/ml was flowed in the chamber for ~3 mins and then the pump was stopped to allow the system to reach equilibrium for 30 mins to an hour. The change in mass on the sensor was visualized by plotting the changes in dissipation (ΔD) as a function of changes in frequency (Δf) for different substrates. The higher the negative frequency shift (−Δf), the higher the adsorbed mass; on the other hand, the higher the dissipation or the slope of the lines, the more viscous or hydrated the molecular films were. After the changes in frequency or dissipation signals reached equilibrium, PBS was circulated again to wash away any excess amount of

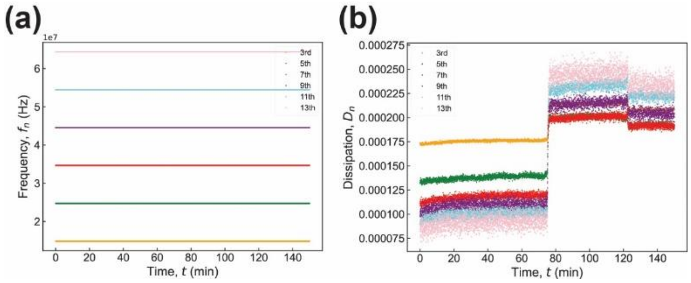

**SI Figure 1 –** Change in mass observed on the sensor via frequency and dissipation vs time.

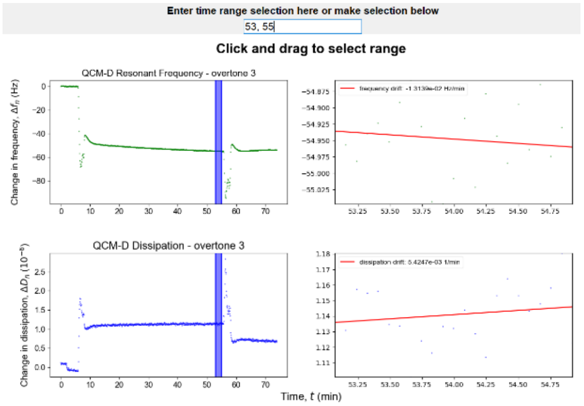


**SI Figure 2.** Before PBS wash for QCM-I analysis.

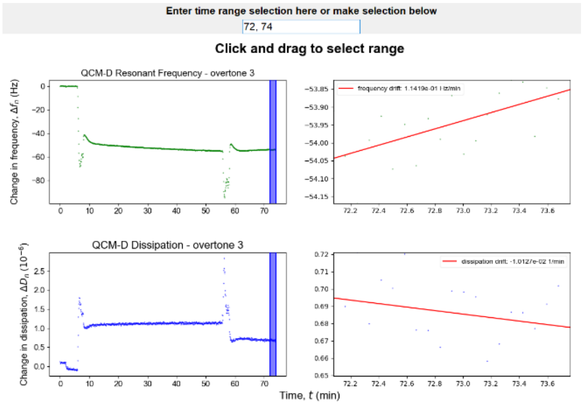


**SI Figure 3.** After PBS wash for QCM-I analysis.

unbound BSA present on the surface. This resulted in a slight reduction in the frequency shift. After waiting for 10-30 mins for the rinsing step to reach equilibrium the experiment and the data collection was stopped.

After every experiment, water and 2% SDS, followed by water, were circulated into the system to clean the inside of the tubing and the connection ports. Then, after purging the chamber of the solutions, the crystal was removed from the chamber. Each measurement was performed at least 3 times independently (N = 3).

The BSA experiment with a QCM-I device described above was replicated in a similar manner using a QSense device in order to test and ensure the consistency of pyQCM-BraTaDio across multiple experimental devices. This experiment is described below.

Prior to use, gold-coated quartz sensors were placed in a 5:1:1 $H_2O/H_2O_2/NH_4O$ mixture at 75°C for 10 minutes and later removed to be rinsed thoroughly with deionized water and dried with nitrogen gas.

Quartz Crystal Microbalance with Dissipation (Q-Sense E1, Biolin Scientific) experiments were performed to determine BSA real-time adsorption behavior by measuring frequency and dissipation shifts. A 5 MHz clean quartz crystal sensor was used to record a stable frequency and dissipation baseline in air. Then the chamber was filled with PBS pH 7.4 buffer and allowed to reach a steady baseline, next switched to the BSA solution at a 1 mg/ml concentration. Finally, PBS is introduced again to wash the excess and measure the adsorbed BSA.

**SI Table 2.** Comparison of results from similar experiments between QCM-I and QSense. Note the fundamental overtone is omitted here due to the inherent noise it possesses. QCM-I selection was made in range t = [3138, 3305] seconds, and QSense in range t = [8725, 8986] seconds.

| | **QCM-I** | **QSense** |
|---|---|---|
| **3$^{rd}$ Overtone Change in Frequency Average, *Δf* (Hz)** | -55.03 | -72.24 |
| **3$^{rd}$ Overtone Change in Dissipation Average, *ΔD* ($10^{-6}$)** | 1.025 | 1.188 |
| **5$^{th}$ Overtone Change in Frequency Average, *Δf* (Hz)** | -87.74 | -116.5 |

| | | |
|---|---|---|
| **5th Overtone Change in Dissipation Average, *ΔD* ($10^{-6}$)** | 1.206 | 1.173 |
| **7th Overtone Change in Frequency Average, *Δf* (Hz)** | -123.3 | -159.3 |
| **7th Overtone Change in Dissipation Average, *ΔD* ($10^{-6}$)** | 1.403 | 1.141 |
| **9th Overtone Change in Frequency Average, *Δf* (Hz)** | -157.8 | -200.7 |
| **9th Overtone Change in Dissipation Average, *ΔD* ($10^{-6}$)** | 1.585 | 1.139 |
| **11th Overtone Change in Frequency Average, *Δf* (Hz)** | -195.8 | -241.5 |
| **11th Overtone Change in Dissipation Average, *ΔD* ($10^{-6}$)** | 1.870 | 1.172 |
| **13th Overtone Change in Frequency Average, *Δf* (Hz)** | -235.8 | -279.9 |
| **13th Overtone Change in Dissipation Average, *ΔD* ($10^{-6}$)** | 2.081 | 1.126 |
| **Sauerbrey Mass $\left(\frac{ng}{cm^2}\right)$** | 324.4 | 372.5 |
| **Crystal Thickness (mm)** | 0.3368 | 0.3371 |
| **Shear Dependent Compliance $\left(\frac{1}{Pa}\right)$** | 0.0204 | 0.0219 |

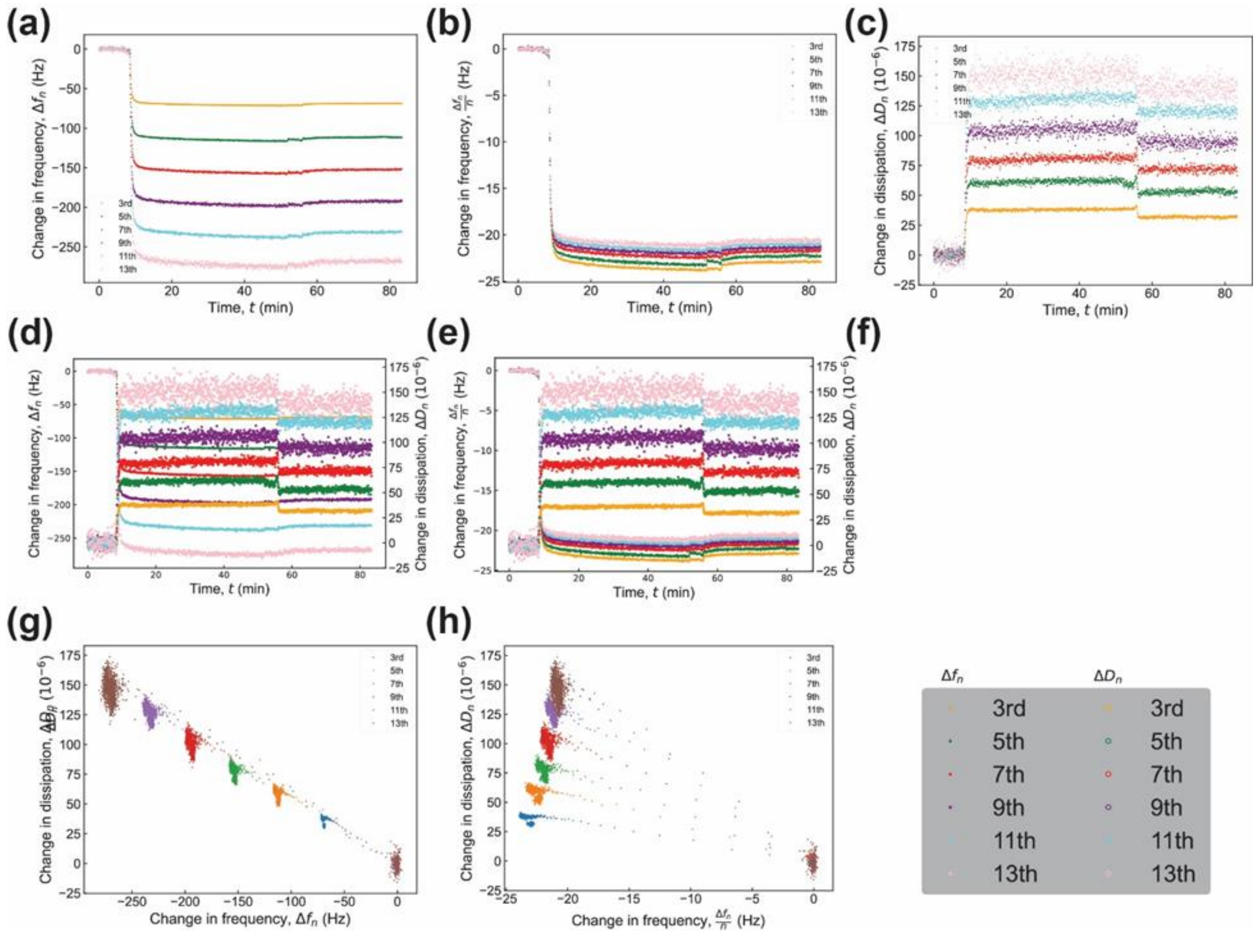


**SI Figure 4. Plots generated by BraTaDio for a film formed from a solution of BSA at 1 mg/mL in PBS adsorbed to an Au-coated quartz crystal.** (a) Change in frequency $\Delta f_n$ as a function of time $t$, (b) change in frequency normalized by overtone order, $\Delta f_n / n$ as a function of time $t$, (c) corresponding change in dissipation $\Delta D_n$ as a function of time, (d) change in frequency normalized by overtone order, $\Delta f_n / n$ and corresponding change in dissipation $\Delta D_n$ as a function of time for $n$ = 5 and 7 for clarity, (e) change in dissipation $\Delta D_n$ as a function of change in frequency normalized by overtone order, $\Delta f_n / n$, for $n$ = 3, 5, and 7 for clarity, and (f) temperature T as a function of time. Data collected with a QSense system.

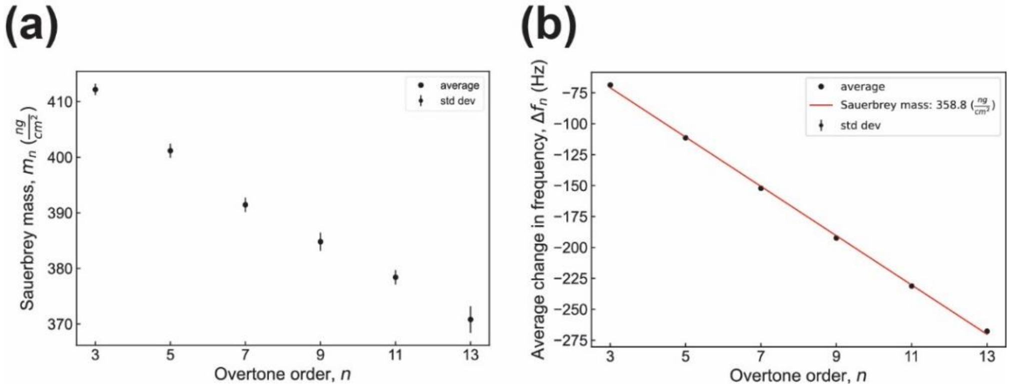


**SI Figure 5. Sauerbrey mass as a function of overtone order.** (a) Sauerbrey mass calculated as a funciton of each individual overtone order and (b) Sauerbrey mass calculated from performing a fit to the average change in frequency as a function of overtone order for the data range shown in SI Figrue 4 (BSA after PBS wash).

### 3 Customize Plot Options

The aspect of the plots can be customized by accessing the *Customize Plots Option* button. The basic customization parameters are font size, font type, color palette, tick direction, time scale for plots in which the data is plotted as a function of time, figure output formats, and plotting ranges. SI Figure 6 shows the various plot customizations, while SI Figure 7 shows two plots of the same data using different plotting options.

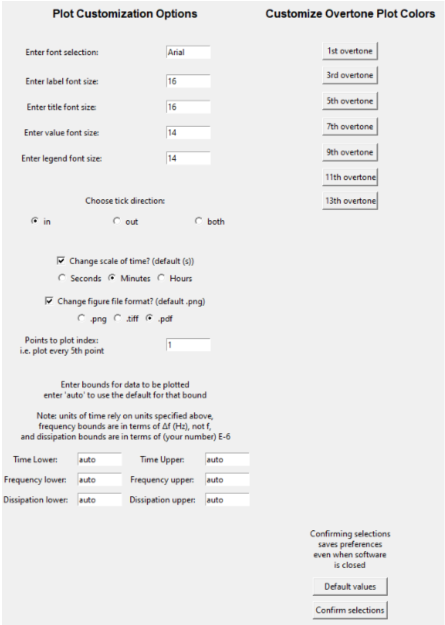


**SI Figure 6.** Plot customization options window

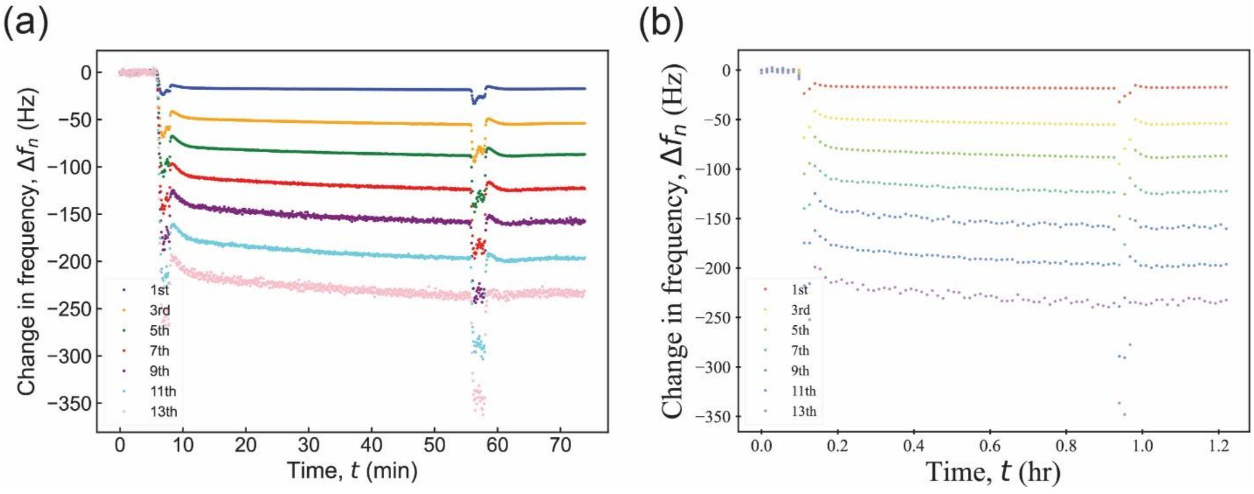


**SI Figure 7.** (a) Example plot using Arial font sizes 16 and 14 for axes titles and values, respectively, plotting every data point with pre-determined color palette. (b) Example plot using Times New Roman font sizes 20 and 12 for axes titles and values, respectively, plotting every 5th data point with a custom color palette.
